# How Older Adults Regulate Lateral Stepping on Narrowing Walking Paths

**DOI:** 10.1101/2023.05.12.540514

**Authors:** Meghan E. Kazanski, Joseph P. Cusumano, Jonathan B. Dingwell

## Abstract

Walking humans often navigate complex, varying walking paths. To reduce falls, we must first determine how older adults purposefully vary their steps in contexts that challenge balance. Here, 20 young (21.7±2.6 yrs) and 18 older (71.6±6.0 yrs) healthy adults walked on virtual paths that slowly narrowed (from 45 cm to as narrow as 5 cm). Participants could switch onto an “easier” path whenever they chose. We applied our Goal Equivalent Manifold framework to quantify how participants adjusted their lateral stepping variability and step-to-step corrections of step width and lateral position as these paths narrowed. We also extracted these characteristics where participants switched paths. As paths narrowed, all participants reduced their lateral stepping variability, but older adults less so. To stay on the narrowing paths, young adults increasingly corrected step-to-step deviations in lateral position more, by correcting step-to-step deviations in step width *less*. Conversely, as older adults also increasingly corrected lateral position deviations, they did so *without* sacrificing correcting step-to-step deviations in step width, presumably to preserve balance. While older adults left the narrowing paths sooner, several of their lateral stepping characteristics remained similar to those of younger adults. While older adults largely maintained overall walking performance *per se*, they did so by changing how they balanced the competing stepping regulation requirements intrinsic to the task: maintaining position vs. step width. Thus, balancing how to achieve multiple concurrent stepping goals while walking provides older adults the flexibility they need to appropriately adapt their stepping on continuously narrowing walking paths.

## INTRODUCTION

Daily walking environments contain complex arrangements of natural and built features, including terrain irregularities (Matthis et al., 2018), and myriad fixed (Twardzik et al., 2019) and moving obstacles (Moussaïd et al., 2011). Such features define the often highly irregular walking paths individuals must navigate in common contexts like crowded sidewalks, or cluttered store aisles. Individuals must continuously enact goal-directed stepping adjustments to remain on these continuously changing paths. As a result, people rarely perform prolonged, straight-ahead walking in daily life (Orendurff et al., 2008).

Despite its limited generalizability to real-world contexts, steady walking on straight invariant paths remains a dominant paradigm for extracting locomotor control principles (Herold et al., 2019; Vistamehr et al., 2016; Wang and Srinivasan, 2014). A critical, but unaddressed, locomotor control problem with direct real-world relevancy is how people enact continuous stepping adjustments on complex, varying walking paths.

It is especially critical to study how older adults enact ongoing stepping adjustments. Older adults experience high rates of falls (Burns and Kakara, 2018) that often result from inadequate stepping adjustments in challenging environments (Li et al., 2006; Rubenstein, 2006). Active modulation of lateral (i.e., side-to-side) stepping is particularly crucial to maintaining balance (Bruijn and van Dieën, 2018) while walking (Dingwell and Cusumano, 2019) along irregular paths or while maneuvering (Desmet et al., 2022). Healthy older adults exhibit generally greater lateral stepping variability during continuous, straight-ahead walking (Owings and Grabiner, 2004; Skiadopoulos et al., 2020), although connecting these changes to fall risk remains challenging (Beauchet et al., 2009; Brach et al., 2005; Moe-Nilssen and Helbostad, 2005). We should instead address how older adults *modulate* their lateral stepping on varying walking paths that better reflect circumstances likely to precipitate falls.

Older adults indeed exhibit increased lateral stepping variability, as compared to younger adults, when traversing irregular walking paths like walking between intermittent obstacles (Maidan et al., 2018) or stepping onto targets (Sun et al., 2017; Zhang et al., 2021). The sources of such age-related increased variability remain unclear. Such age-related changes in lateral stepping variability may reflect detrimental increases in physiological noise (Christou, 2011) that reduce lateral stepping accuracy (Kuo and Donelan, 2010) and compromise locomotor stability (Roos and Dingwell, 2010). Alternatively, age-related altered variability may reflect beneficial neuromuscular compensation (Eckardt and Rosenblatt, 2018) that allows older adults to exhibit considerable variability in select movement variables (Todorov, 2004) while still reliably accomplishing the goal-directed walking (Dingwell et al., 2017) necessary in challenging environments (Kazanski et al., 2020). Before determining how age-related stepping variability relates to falling risk in older adults, we must first determine the extent to which lateral stepping variability arises from which underlying sources (e.g., noise, motor redundancy, or changing task goals), especially on varying walking paths that require purposeful, goal-directed stepping adjustments.

We previously identified how stepping variability arises from goal-directed stepping regulation processes using an analytical framework entailing Goal Equivalent Manifolds (GEMs) (Cusumano and Dingwell, 2013; Dingwell et al., 2010). The GEM framework is based on formulating *goal functions* that pose testable hypotheses on how humans regulate stepping movements to achieve different walking tasks (Dingwell et al., 2010). The GEM is the set of all combinations of step-to-step observables (e.g. foot placements) that equally achieve some hypothesized walking task goal like “maintain speed” (Dingwell et al., 2010) or “maintain step width” (Dingwell and Cusumano, 2019), etc. We previously used the GEM framework to demonstrate that versatile walking is facilitated by inherently *multi*-objective lateral stepping regulation: humans can trade-off between competing task goals to maintain either step width or lateral body position (Dingwell and Cusumano, 2019). During continuous, straight walking, healthy adults prioritize minimizing errors from a target step width, while to a much lesser extent correcting errors in their lateral body position from the center of their walking path (Dingwell and Cusumano, 2019). Humans adapt these lateral stepping regulation processes to different contexts by re-weighting prioritization of these step width vs. lateral body position goals (Desmet et al., 2022; Render et al., 2021) and/or increasing correction of errors in *both* variables (Dingwell et al., 2020; Kazanski et al., 2020). Healthy aging does *not* alter lateral stepping regulation during straight or laterally-perturbed walking (Kazanski et al., 2020). We expect that these same types of GEM analyses should allow us to discern how age affects the lateral stepping regulation that people enact when walking on varying paths.

Here, we presented healthy young and older adults with systematically-narrowing virtual walking paths. We expected this paradigm would prompt participants to continuously adjust their lateral stepping regulation to maintain both step widths *and* lateral body positions on these narrowing paths. We predicted all (young and older) participants would reduce their variability as paths narrowed and would progressively increase emphasis given to regulating body position, trading-off step width regulation as paths narrowed. With respect to age, we expected the older adults would exhibit greater variability (Lowry et al., 2017) than the younger, but would also retain greater prioritization of step width to preserve lateral balance. Noting that some older adults decline to engage with sufficiently narrow paths at all (Butler et al., 2015), we allowed participants to leave the narrowing path at any time. We then quantified whether groups exhibited different lateral stepping regulation when participants left the narrowing path.

## METHODS

### Participants

We analyzed data from the same cohort of 20 young healthy (YH) and 18 older healthy (OH) adults as in (Kazanski and Dingwell, 2021) (Table 1). We consented and screened all participants prior to study participation using protocols approved by the IRB at Pennsylvania State University.

**Table 1.** Young Healthy and Older Healthy physical characteristics and assessment scores. Functional balance ability was assessed using Timed Up and Go (TUG) and Four Square Step Test (FSST). Balance confidence was assessed using the Activities-Specific Balance Confidence (ABC) Scale and Iconographical Falls Efficacy Scale (I-FES). All values except Sex are given as Mean ± Standard Deviation. Statistical p-values are from two-sample t-test or Mann-Whitney U tests, as relevant.

| **Characteristic:** | **Young Healthy (YH):** | **Older Healthy (OH):** | **p-value** |
| --- | --- | --- | --- |
| Sex [M/F] | 9 / 11 | 5 / 13 |  |
| Age [yrs] | 21.7 ± 2.6 | 71.6 ± 6.0 | **< 0.001** |
| Body Height [m] | 1.73 ± 0.08 | 1.70 ± 0.10 | 0.305 |
| Body Mass [kg] | 69.9 ± 12.1 | 70.0 ± 10.5 | 0.983 |
| **Assessment:** |  |  |  |
| TUG [s] | 6.62 ± 0.75 | 8.20 ± 1.92 | **0.004** |
| FSST [s] | 7.63 ± 1.37 | 9.19 ± 1.53 | **0.002** |
| ABC Score [/100 %] | 96.8 ± 2.5 | 93.1 ± 5.5 | **0.023** |
| I-FES [/40] | 12.3 ± 1.9 | 16.1 ± 3.9 | **0.001** |

[Insert Table 1 about here]

### Protocol and Data Collection

Participants performed five, 4-minute trials walking on a series of virtual paths projected onto a 1.2m wide treadmill (Motek M-Gait system, Amsterdam, Netherlands), as detailed in (Kazanski and Dingwell, 2021). Each trial consisted of eight path-walking bouts. Each bout presented two parallel, 25m-long virtual paths projected onto the left and right treadmill belts (Fig. 1A). One path gradually narrowed from an initial width of 0.45m to a final width of 0.05m according to:

**Figure 1.**
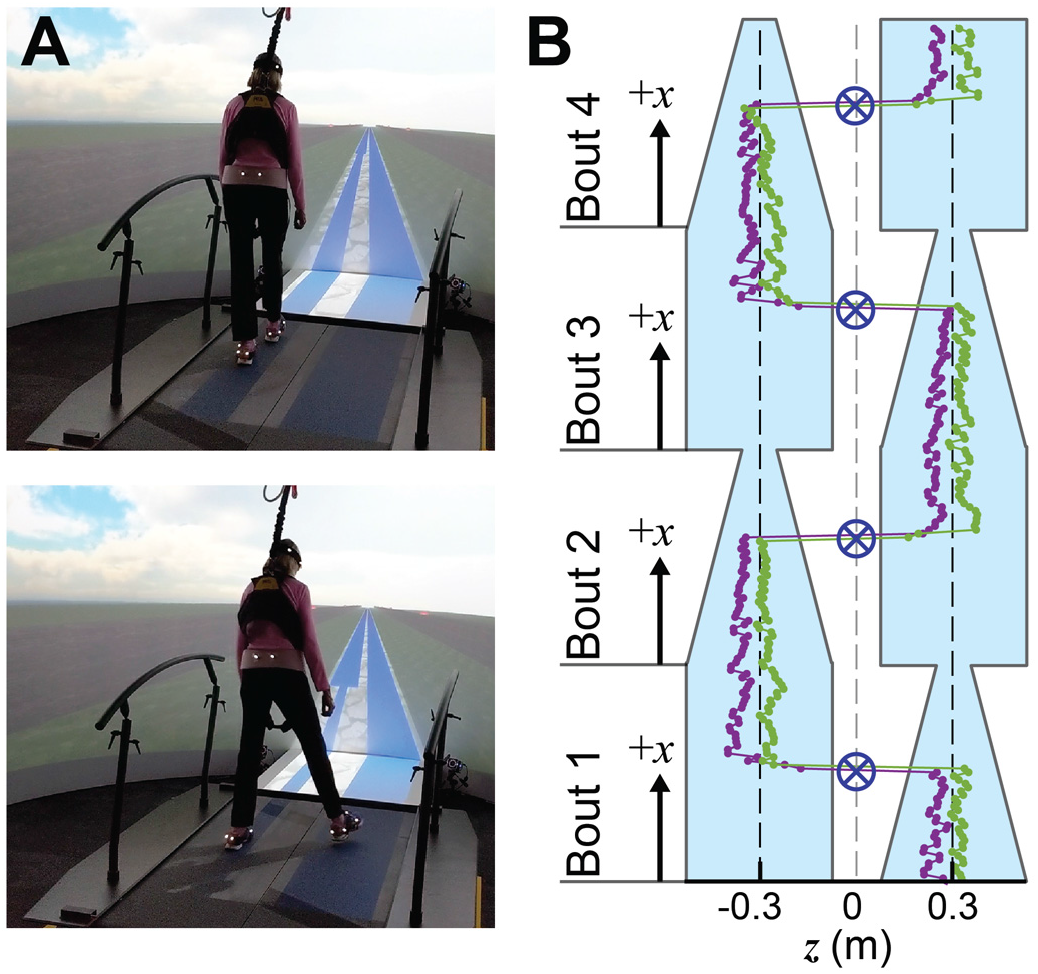
**(A)** For each walking bout, participants walked on a continuously-narrowing virtual path in an “M-Gait” system (Motek, Amsterdam, Netherlands). Participants were instructed to remain on the path as long as possible, while minimizing stepping errors outside of the path bounds. The narrowing path width gradually decreased from 0.45m to 0.05m. Meanwhile, an adjacent path remained fixed at 0.45m. Participants could switch off the narrowing path to the adjacent path at any time. **(B)** Each trial consisted of eight walking bouts. Narrowing vs. fixed-width paths alternated sides of the treadmill every 25m, at which point a new walking bout started and the path progression (*x*) was reset. Participants always switched to the fixed-width path before each new walking bout started. Example series of lateral (*z*-direction) left and right foot placements are shown as purple and green markers, respectively. Path switches (blue crosshairs) occurred when the lateral CoM trajectory crossed the centerline of the treadmill (dashed grey line). Dashed black lines indicate walking path centers.

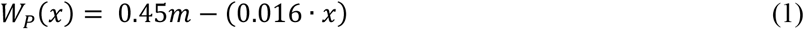

where *W*_*P*_(*x*) is the narrowing path width and *x* is the walker’s path progression, which increased from 0m to 25m during each bout. Meanwhile, the adjacent path on the opposite treadmill belt remained fixed at 0.45m wide. A new walking bout began every 25m, at which point the narrowing and fixed-width paths alternated sides of the treadmill (Fig. 1B).

This task required ever-increasing stepping accuracy as the path narrowed, likely imposing a speed-accuracy trade-off (Roerdink et al., 2021). We therefore fixed treadmill speed at 0.75 m/s. This speed is within the range older adults exhibit when asked to walk with narrow-base gait (Schrager et al., 2008). We held speed fixed to preclude the additional confound of differences in self-selected speeds. The optic flow speed of the virtual environment was matched to this treadmill speed.

As people walk along *any* path, they adjust each foot placement to trade off maintaining step width (to stay balanced) for lateral position (to stay on their path) (see Supplement #1) (Dingwell and Cusumano, 2019; Kazanski et al., 2020). Thus, as paths presented here narrowed, this task slowly but increasingly constrained where people could place their feet to negotiate this trade-off. This task directly challenged each participant’s ability to balance those competing task goals of maintaining balance while staying on their path.

Participants began each bout on the narrowing path and were instructed to remain on it as long as possible, while minimizing stepping errors off of the path. Participants chose when to switch over to the adjacent fixed-width path. Participants completed a 60s wash-out trial of steady-state walking after each path walking trial.

Participants rested (≥ 2-min) between all trials.

Kinematic data were collected as described in (Kazanski and Dingwell, 2021). Briefly, participants wore 16 retroreflective markers. Only left and right heel and posterior pelvis markers were used for the present analyses. Marker trajectories were collected at 120 Hz using a 10-camera motion capture system (Oxford Metrics, Oxford, UK), then post-processed in Vicon Nexus and exported to MATLAB (MathWorks, Natick, MA).

### Relevant Stepping Variables

As we walk, we must choose, at each new step, where to place each foot relative to our path. (Dingwell and Cusumano, 2019). Here, we designate {*z*_*Ln*_, *z*_*Rn*_} to indicate the lateral (*z*-direction) left and right foot placements relative to the path center (Fig. 2A) at each step, *n*. Note, we define body position (*z*_*Bn*_) as the mid-point between {*z*_*Ln*_, *z*_*Rn*_}, as this yields an unambiguous once-per-step estimate of the lateral center-of-mass location at each step (Desmet et al., 2022; Dingwell and Cusumano, 2019).

**Figure 2.**
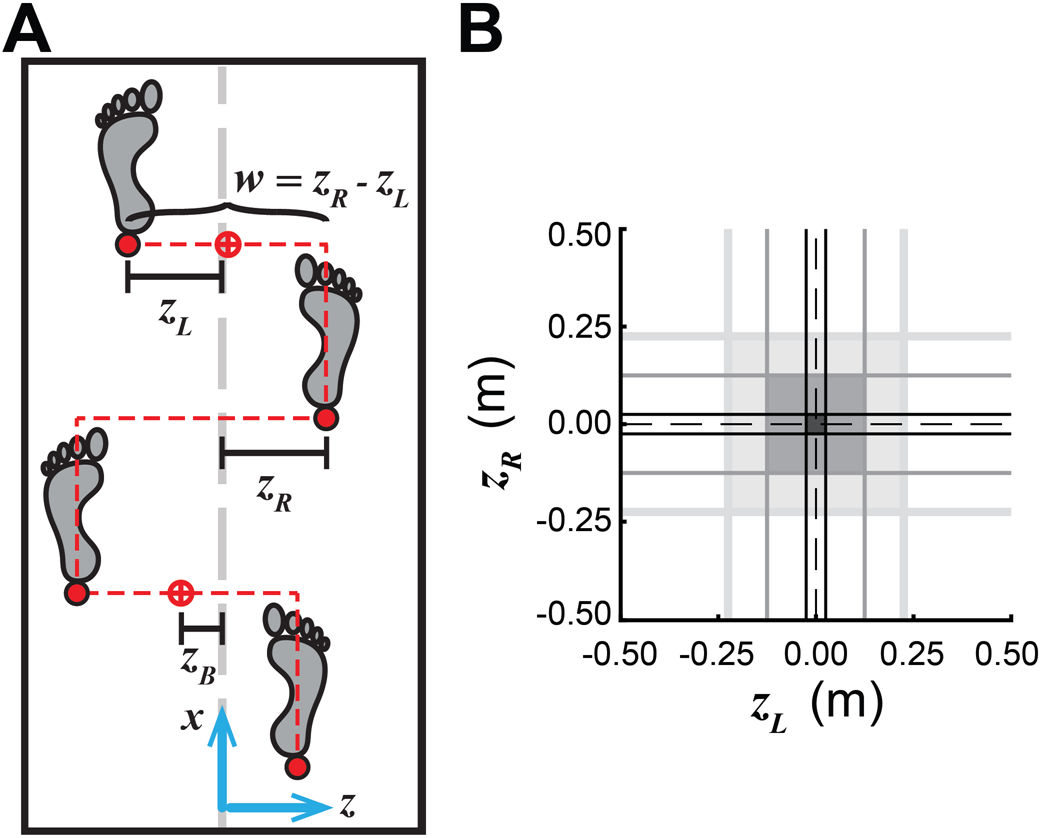
**(A)** Walking humans coordinate left and right foot placements from step-to-step to regulate lateral body position (*z*_*B*_) relative to path center and step width (*w*). Each step is defined by both foot placements (*z*_*L*_, *z*_*R*_) associated with that step. Lateral body position is the midpoint between them: *z*_*B*_ = ½(*z*_*L*_ + *z*_*R*_). Step width is the difference between them: *w* = (*z*_*R*_ − *z*_*L*_) (Dingwell and Cusumano, 2019). **(B)** At any *W*_*P*_(*x*_*n*_), successful left and right foot placements will land within the path bounds (Eq. 1), represented as a square region in the [*z*_*L*_, *z*_*R*_] plane, centered at (0,0) with area [*W*_*P*_(*x*_*n*_)]^2^. As *W*_*P*_(*x*_*n*_) decreases along the path, the region of viable foot placements also decreases from wide (light grey square), to intermediate (darker grey square), to narrow (darkest grey square).

We previously showed (Dingwell and Cusumano, 2019) that when humans walk on paths with constant width, they do *not* directly regulate {*z*_*Ln*_, *z*_*Rn*_}. Instead, they coordinate {*z*_*Ln*_, *z*_*Rn*_} to multi-objectively maintain some combination of step width (*w*_*n*_) and lateral body position (*z*_*Bn*_) relative to their path. Foot placements, {*z*_*Ln*_, *z*_*Rn*_}, are directly mathematically related to these goal states, {*z*_*Bn*_, *w*_*n*_}, as described in Supplement #1.

For each walking bout, we extracted step-to-step time series of {*z*_*Ln*_, *z*_*Rn*_} using the lateral positions of respective left and right heel markers at each consecutive step *n* ( {1, …, *N*} (Zeni et al., 2008). We then computed time series of {*z*_*Bn*_, *w*_*n*_} from these {*z*_*Ln*_, *z*_*Rn*_} (see Supplement #1). We then also extracted corresponding step-to-step time series of *W*_*P*_(*x*_*n*_) for each walking bout.

We analyzed only steps taken along the narrowing path, until the second-to-last step before path switching. Each path switch instance occurred when the pelvic centroid crossed the treadmill midline. We extracted the narrowing path width ((*W*_*P*_)_*switch*_) at each path switch instance.

We first pooled all time series for all subjects by group (YH or OH). Then, we separated these stepping data into overlapping “bins” that covered the range of gradually-decreasing path width (*W*_*P*_), as in (Kazanski and Dingwell, 2021). We required at least two *consecutive* steps from any given trial in each bin; single steps were discarded. Sixty-six total overlapping bins each contained all steps that occurred over a 0.075m range of *W*_*P*_, with new bins assigned every 0.005m of *W*_*P*_. Each bin’s designated width, *W*_*P*_, was taken as the mid-point of possible *W*_*P*_ for that bin. This allowed us to characterize how lateral stepping changed along the narrowing path (e.g., Fig. 3A). For these pooled group data within each of the 66 bins, we then sub-sampled 50 data sets by extracting a random 95% of the data in that bin.

**Figure 3.**
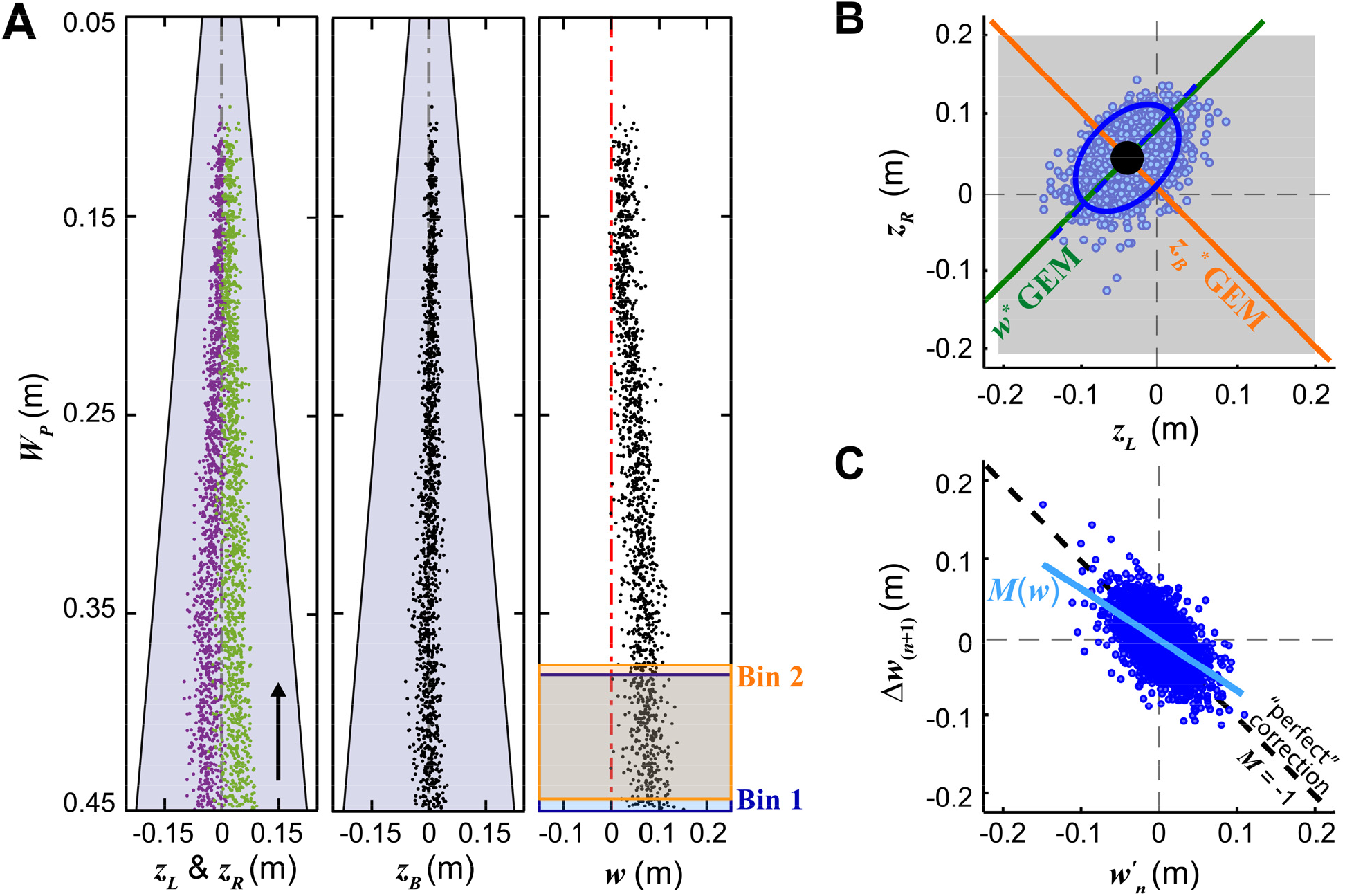
**(A)** Lateral stepping time-series data are shown for a single OH participant, relative to corresponding path width (*W*_*P*_). Each marker represents a single step. Variables relative to path center include: left (*z*_*L*_; purple) and right (*z*_*R*_; green) foot placements (left), lateral body position (*z*_*B*_; middle) and step width (*w*; right). On the {*z*_*L*_, *z*_*R*_} and *z*_*B*_ plots, vertical grey dashed lines indicate path center. On the *w* plot, the red vertical dashed line indicates *w* = 0. Overlayed on the *w* plot, two adjacent, overlapping bins are shown to demonstrate the binning procedure. Bin 1, shown in blue and centered at *W*_*P*_ = 0.4125m, contains all steps taken on path widths between 0.45m and 0.375m. Bin 2, shown in orange and centered at *W*_*P*_ = 0.4075m, contains all steps taken on path widths between 0.445m and 0.370m. Corresponding bins spanned the length of the narrowing path to capture all enacted foot placements. For subsequent analyses, such lateral stepping time-series data were pooled separately for each YH and OH age group across walking bouts, trials and participants. **(B)** Group-wise lateral foot placements plotted in [*z*_*L*_, *z*_*R*_] at a given *W*_*P*._ Individual steps are shown as single markers. Lateral stepping regulation variables [*z*_*B*_, *w*] map onto [*z*_*L*_, *z*_*R*_] as mutually-orthogonal GEMs representing constant lateral body position (*z*_*B*_ ^*^= constant; orange) and constant step width (*w*^*^= constant; green). The fitted 95% prediction ellipse (blue) is centered at the mean operating point (*z*_*L*_ ^*\**^ *z*_*R*_ ^*\**^) (black dot). The major ellipse axis is represented by the blue dashed line. **(C)** Group-wise {*wʹ*_*n*_, Δ*w*_(n*+1*)_} demonstrate deviations from *w*^*\**^ at step *n* (*wʹ*_*n*_ = *w*_*n*_-*w*^*^) and corrections on subsequent step *n*+1 (Δ*w*_(*n*+1)_ = *w*_(n*+1*)_-*w*_*n*_) at a given *W*_*P*_. Individual stepping combinations are represented by single markers. The diagonal solid blue line indicates the linear error correction, with slope *M*(*w*) computed via least-squares fits that defines the extent of *w* correction in this sub-sample. The diagonal dashed black line indicates “perfect” error correction: i.e., *M*(*w*) = −1. These analyses were applied within all sub-samples across all quasi-constant *W*_*P*_. Identical analyses were applied to group-wise [z_*B*_*ʹ*_*n*_, Δz_*B*(n*+1*)_].

### Stepping Variability

To characterize each group’s stepping behavior within each bin, we first visualize how steps {*z*_*Ln*_, *z*_*Rn*_} directly relate to goal states {*z*_*Bn*_, *w*_*n*_} by plotting them in the [*z*_*L*_, *z*_*R*_] plane (Fig. 3B) (Desmet et al., 2022; Dingwell and Cusumano, 2019). In this [*z*_*L*_, *z*_*R*_] plane, orthogonal diagonal lines define Goal Equivalent Manifolds (Cusumano and Cesari, 2006) for maintaining either constant *z*_*B*_ or constant *w* (see Supplement #1). The distribution of lateral stepping variability within the [z_*L*_, *z*_*R*_] plane will this reflect how strongly people prioritize *z*_*B*_ vs. *w* regulation (Supplement #1).

We first plotted all group-wise lateral steps in [z_*L*_, *z*_*R*_] for each sub-sample. We fit 95% prediction ellipses to these stepping distributions, defined by scaling the eigenvalues of the {*z*_*L*_, *z*_*R*_} covariance matrices by the 95^th^ percentile critical value of the Chi-Square Distribution (*χ*^2^(0.05) = 0.5991) (Desmet et al., 2022). Each such ellipse can then be fully defined by its area, aspects ratio (a measure of “shape”), and orientation (Supplement #1; Fig. S1-1).

We computed the relative area (*A*_*Rel*_) of each variability ellipse relative to its square path area:

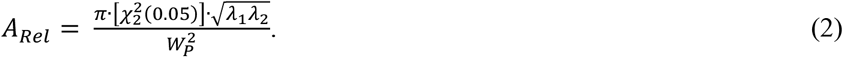

A larger *A*_*Rel*_ indicates participants varied their lateral steps over a greater proportion of their available path width, *W*_*P*_. We computed the aspect ratio of each ellipse as the ratio of covariance matrix eigenvalues (*λ*_1_/*λ*_2_), where *λ*_1_/*λ*_2_ >> 1 indicates strong elongation along the direction of the ellipse major axis. We computed the orientation (Δθ) of each ellipse as the angular deviation of the ellipse major axis from the +*w*^*\**^ GEM (Supplement #1; Fig. S1-1).

We then averaged all 50 sub-sample values of each feature within each bin. We plotted these average *A*_*Rel*_, *λ*_1_/*λ*_2_, and Δθ across *W*_*P*_ for each group: YH and OH. We expected both groups would (necessarily) increase their *A*_*Rel*_ as *W*_*P*_ decreased. However, we expected OH would exhibit consistently more-variable (i.e., larger *A*_*Rel*_) steps across the range of *W*_*P*_. We further expected both groups would increase their prioritization of *z*_*B*_ over *w* regulation as *W*_*P*_ narrowed, yielding lower *λ*_1_/*λ*_2_ and/or higher |Δθ| with decreasing *W*_*P*_. However, we expected OH would exhibit consistently greater prioritization of *w* (thus higher *λ*_1_/*λ*_2_, lower |Δθ|) relative to YH across the range of *W*_*P*_.

### Error Correction

The above measures do not quantify the extent to which walking people correct errors in *w* and/or *z*_*B*_ from each step to the next (Dingwell and Cusumano, 2015, 2019). Participants attempting to achieve some target *w*^*\**^ at each step will exhibit small deviations on any *n*^*th*^ step: *w*_*n*_*’* = *w*_*n*_ - *w*^*\**^. Such deviations might then be corrected on the next step: Δ*w*_*n+1*_ = *w*_*n+1*_ – *w*_*n*_. For each sub-sample in each bin, we plotted all combinations of Δ*w*_*n+1*_ vs. *w*_*n*_*’* (Fig. 3C). We computed linear slopes *M*(*w*) of each plot to quantify the extent of error correction, where *M*(*w*) = −1 indicates “perfect” correction, *M*(*w*) < −1 over-correction, and *M*(*w*) > −1 under-correction (Dingwell and Cusumano, 2015). We used this same procedure to quantify *z*_*B*_ error correction: *M*(*z*_*B*_). We averaged all 50 sub-sample values of each *M*(*z*_*B*_) and *M*(*w*) within each bin and plotted average *M*(*z*_*B*_) and *M*(*w*) across *W*_*P*_ for each group.

We expected that as *W*_*P*_ decreased, both groups would increase correction of *z*_*B*_ (i.e., *M*(*z*_*B*_) → −1) by trading off correction of *w* (i.e., *M*(*w*) → 0). We further expected that, relative to YH, OH would demonstrate stronger prioritization of *w* (i.e., *M*(*w*)_OH_ < *M*(*w*)_YH_) and weaker prioritization of *z*_*B*_ (i.e., *M*(*z*_*B*_)_OH_ > *M*(*z*_*B*_)_YH_).

### When Participants Switched Paths

We also characterized each participant’s lateral stepping at the mean instants they switched off of the narrowing path. We first selected the group-wise bin nearest to each participant’s average (*W*_*P*_)_*switch*_. We then used the groupaverage stepping variability {(*A*_*Rel*_)_*switch*_, (*λ*_1_/*λ*_2_)_*switch*_, (Δ*θ*)_*switch*_)} and error correction {*M*(*z*_*B*_)_*switch*_, *M*(*w*)_*switch*_} at that bin to represent that participant’s lateral stepping at their average path switch. We compared these statistically in Minitab (Minitab, Inc., State College, PA) using two-sample t-tests or Mann-Whitney U tests to assess age group differences for each measure. We used these comparisons to determine if differences in OH lateral stepping regulation might explain why they switched off the narrowing path sooner, at wider (*W*_*P*_)_*switch*_ (Kazanski and Dingwell, 2021).

## RESULTS

Most OH adults switched walking paths sooner at wider path widths, (*W*_*P*_)_*switch*_, than YH (Fig. 4) (Kazanski and Dingwell, 2021).

**Figure 4.**
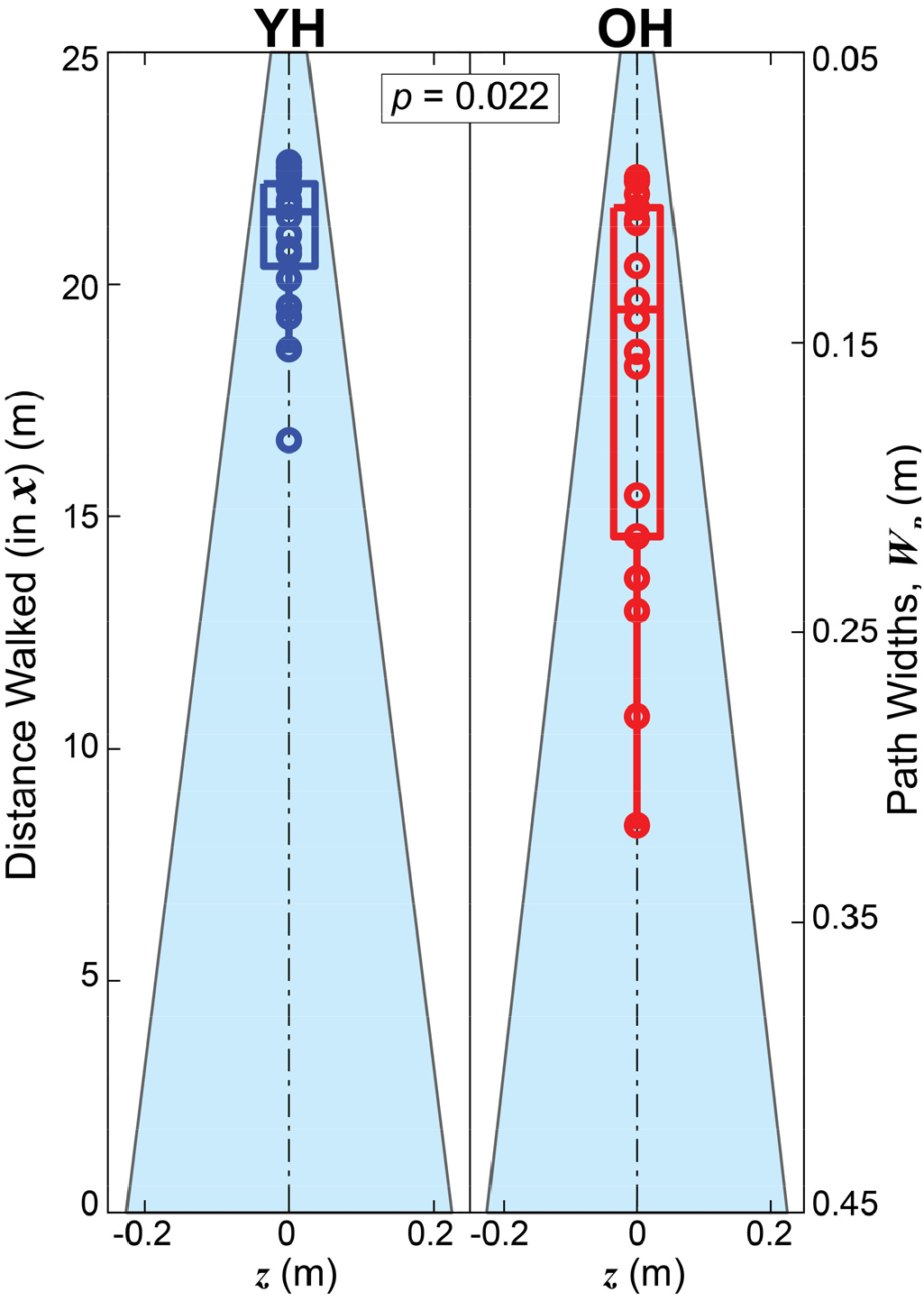
Points along the narrowing paths at which younger (YH; left) and older (OH; right) participants transitioned off of the path. Shaded isosceles trapezoids represent the narrowing paths walked on. The left side axis labels give the distances traveled along the path (i.e., in the +***x*** direction). The right side axis labels give path widths as defined by Eq. (1). Individual average transition points are shown as circles for each participant (YH blue; OH red). Box plots indicate medians, 1^st^ and 3^rd^ quartiles, and whiskers extending to 1.5× interquartile range. OH participants switched walking paths sooner, at wider path widths (p = 0.022; Kazanski et al., 2021).

As paths narrowed, both groups remained within the path bounds (*A*_*Rel*_ < 1), but progressively varied their lateral steps across more of the available *W*_*P*_ (Fig. 5A) (see also Supplement #2). OH consistently varied their lateral steps over a greater proportion of *W*_*P*_ (i.e., larger *A*_*Rel*_) than YH.

**Figure 5.**
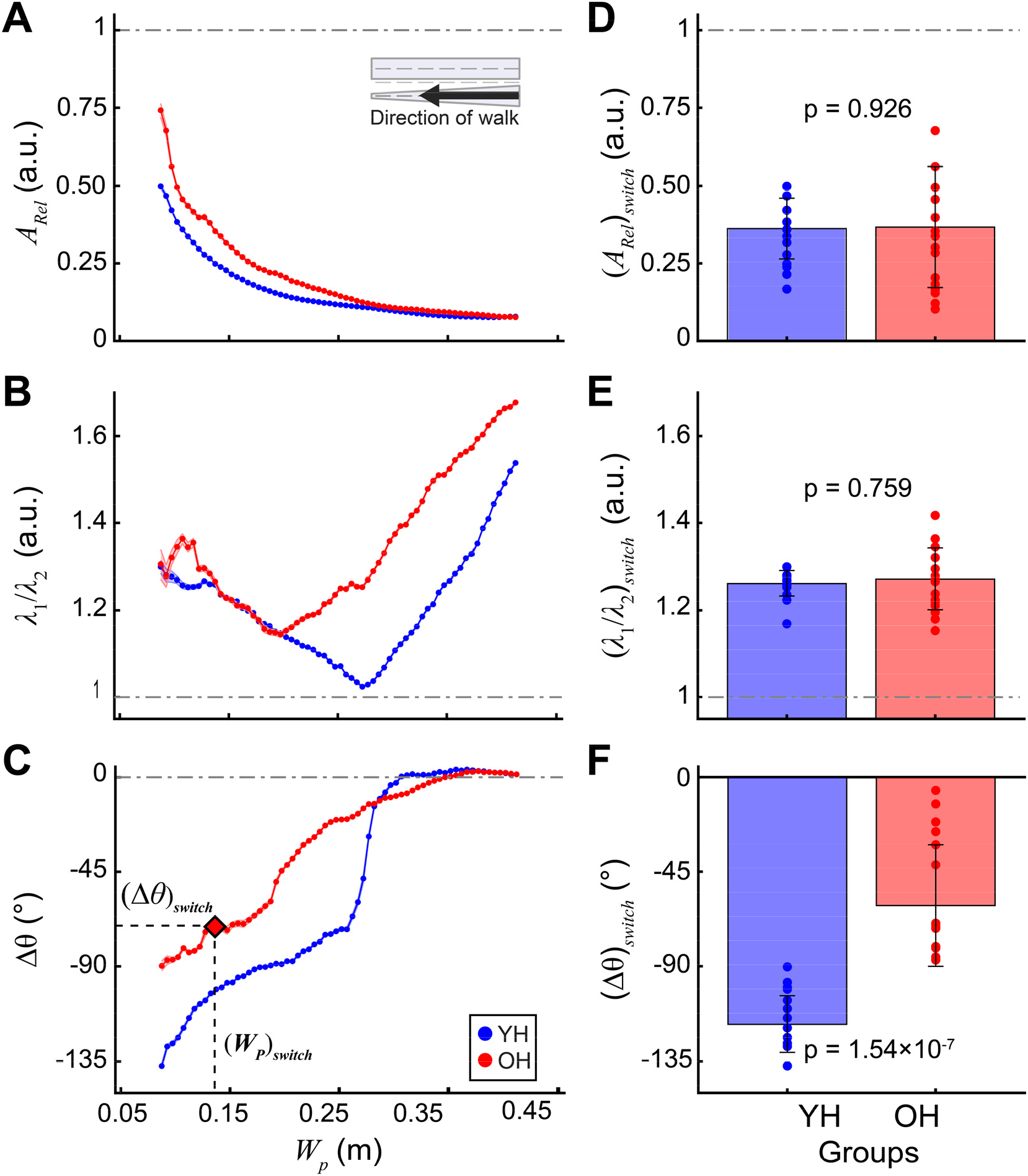
95% prediction ellipses (Figs. 3B, 4) characterized lateral stepping variability in [*z*_*L*_, *z*_*R*_] across bins of decreasing path widths (*W*_*P*_). **(A)** Relative area (*A*_*Rel*_), **(B)** aspect ratio (*λ*_1_/*λ*_2_), and **(C)** orientation (Δθ), each averaged across all sub-samples within each bin (see Fig. 2). Shaded error regions (small, but plotted on all curves) represent ±1 SD between sub-samples. The path width bin at the average instant each participant switched paths, (*W*_*P*_)_*switch*_, was mapped onto each of these plots, as shown in (**C**), to characterize their variability thresholds at path switch: i.e., (*A*_*Rel*_)_*switch*_, (*λ*_1_/*λ*_2_)_*switch*_, and (Δθ)_*switch*_. OH exhibited **(D)** similar (*A*_*RelARel*_)_*switch*_, (YH: 0.36±0.10; OH: 0.37×±0.20; p = 0.926) and **(E)** similar (*λ*_1_/*λ*_2_)_*switch*_ (YH: 1.26±0.03; OH: 1.27±0.07; p = 0.759), but **(F)** less-negative (Δθ)_*switch*_ (YH: −117.38±13.33°; OH: −61.12±28.93°; p = 1.54×10^-7^) relative to YH.

Initially at wider *W*_*P*_, both YH and OH variability was strongly aligned with the step width (*w*^*\**^) GEM (*λ*_1_/*λ*_2_ >> 1 and Δθ ≈ 0°; Fig. 5B-C), consistent with straight-ahead walking (Dingwell and Cusumano, 2019). As paths narrowed, both groups progressively reduced alignment with the step width GEM *toward* alignment with the position (*z*_*B*_^*\**^) GEM. This was indicated by decreasing Δθ (Fig. 5C) and by *λ*_1_/*λ*_2_ that initially decreased, but then increased (Fig. 5B) as Δθ decreased further. Relative to YH, OH adults’ lateral stepping variability deviated less from the step width GEM (Fig. 5C).

Initially, at the widest path widths (*W*_*P*_ → 0.45m), both YH and OH groups exhibited stronger correction of step width errors (*M*(*w*) → −1; Fig. 6B) and weaker correction of position errors (*M*(*z*_*B*_) → 0; Fig. 6A), also consistent with straight-ahead walking (Dingwell and Cusumano, 2019). As paths narrowed (i.e., as *W*_*P*_ decreased), both groups decreased *w* error correction (less negative *M*(*w*); Fig. 6B) while increasing *z*_*B*_ error correction (more negative *M*(*z*_*B*_); Fig. 6A). Relative to YH, OH generally prioritized *w* correction more (*M*(*w*)_OH_ < *M*(*w*)_YH_) and *z*_*B*_ correction less (*M*(*z*_*B*_)_OH_ > *M*(*z*_*B*_)_YH_) throughout (Fig. 6A-B).

**Figure 6.**
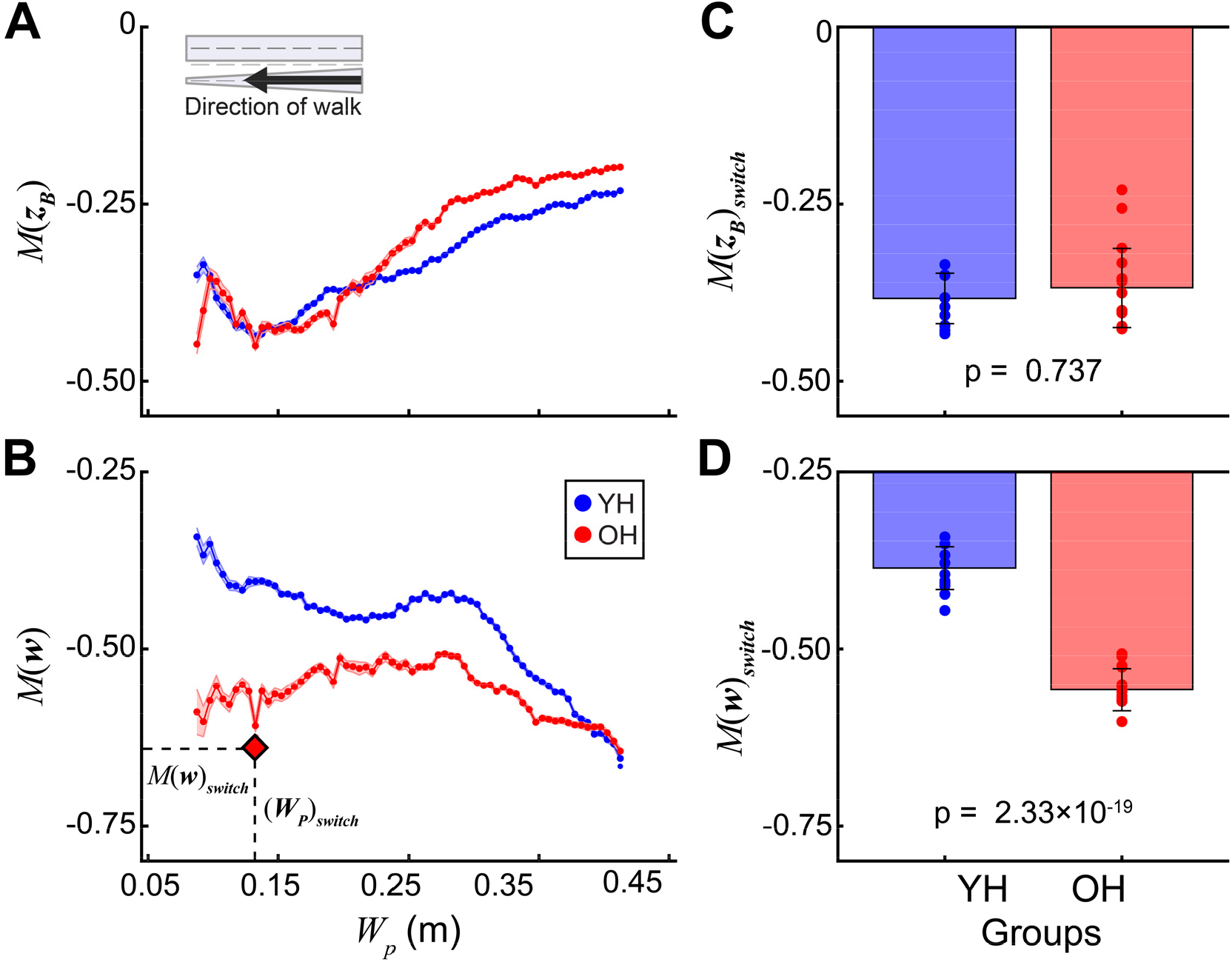
Average of sub-sampled linear error correction slopes for **(A)** lateral body position (*M*(*z*_*B*_)) and **(B)** step width (*M*(*w*) across bins of decreasing path widths (*W*_*P*_). Shaded error regions (small, but plotted on all curves) represent ±1 SD between sub-samples. The path width bin at each participant’s average instant of path switch, (*W*_*P*_)_*switch*_, was mapped onto each of these plots to determine error correction thresholds *M*(*z*_*B*_)_*switch*_ and *M*(*w*)_*switch*_, as shown in **(B)**. OH exhibited **(C)** similar *M*(*z*_*B*_)_*switch*_ (YH: −0.38±0.03; OH: −0.37±0.06; p = 0.737) and **(D)** more-negative *M*(*w*)_*switch*_ (YH: −0.39±0.03; OH: −0.56±0.03; p =2.33×10^-19^) relative to YH.

At the respective path width at which individual participants chose to switch paths, (*W*_*P*_)_*switch*_, YH and OH participants exhibited similar (*A*_*Rel*_)_*switch*_ (Fig. 5D) and (*λ*_1_/*λ*_2_)_*switch*_ (Fig. 5E), but significantly smaller (closer to 0°) (Δθ)_*switch*_ (Fig. 5F). Relative to YH, when OH switched off of the narrowing path, they prioritized position correction (*M*(*z*_*B*_)_*switch*_) similarly (Fig. 6C), but step width correction (*M*(*w*)_*switch*_) more (Fig. 6C).

## DISCUSSION

Humans must continuously navigate walking paths defined by features of their daily environments (Matthis et al., 2018; Twardzik et al., 2019). They do this by exploiting the multi-objective nature of lateral stepping regulation (Dingwell and Cusumano, 2019) to carry out goal-directed stepping adjustments across multiple challenging contexts (Desmet et al., 2022; Dingwell et al., 2020; Kazanski et al., 2020; Render et al., 2021). Any age-related changes in one’s ability to regulate stepping may become particularly relevant when older adults walk in challenging environments. Here, young and older adults walked on continuously-narrowing virtual paths. We applied our Goal Equivalent Manifold (GEM) framework for lateral stepping to quantify how healthy aging alters the structure of lateral stepping variability and stepping regulation processes on these narrowing walking paths.

Prior studies examined how aging (Arvin et al., 2016) and redundancy (Rosenblatt et al., 2014) contribute to altered stepping variability on different paths of constant width. This present study differed: here, the path presented narrowed continuously and never imposed exact foot placements, step widths, or lateral body positions. Participants could thereby execute any of an infinite number of {*z*_*L*_, *z*_*R*_} combinations (within the range of the current path width) to remain on the path. To remain on these paths, all participants progressively reduced their net amount of lateral stepping variability, while filling increasing portions of path width (Fig. 5A, left).

Furthermore, both groups traded-off correcting step width errors to more strongly prioritize correcting lateral position errors (Fig. 6A-B left). In doing so, they thus executed steps that were distributed away from step width GEM alignment toward the position GEM (Fig. 5B-C, left). Thus, both age groups progressively and substantially deviated from the goal-directed stepping characteristics observed during continuous, straight-ahead walking in a manner consistent with our theoretical framework (Dingwell and Cusumano, 2019). This present study further demonstrates how walking humans use the flexibility afforded by multi-objective regulation processes to achieve highly adaptive lateral stepping in diverse contexts (Desmet et al., 2022; Dingwell and Cusumano, 2019; Dingwell et al., 2020; Kazanski et al., 2020; Render et al., 2021).

Healthy aging did not degrade these older adults’ capacities to keep foot placements within the path bounds prior to their switching paths. However, these older adults did consistently exhibit greater relative variability within the path bounds compared to young adults (Fig. 5A, left). This is consistent with prior findings from fixed-width narrow paths that also showed older adults generally did not reduce lateral stepping variability to the same extent as younger adults (Arvin et al., 2016; Schrager et al., 2008).

This study is the first, to our knowledge, to quantify how healthy aging affects not only the amount and structure of lateral stepping variability, but also step-to-step regulation, in a continuously-varying context. Older adults differed in that they structured their lateral stepping to be consistently more closely aligned to the step width GEM (Fig. 5B-C, left). This corresponded with their consistently stronger step width correction (Fig. 6B, left) and weaker lateral position correction (Fig. 6A, left) relative to young adults. This increased prioritization of *w* regulation is consistent with findings that older adults tend to rely on wider, more variable lateral foot placements to maintain balance while walking (Hollman et al., 2011; Owings and Grabiner, 2004). Thus, our work shows healthy older adults also exploit multi-objective lateral stepping regulation to re-weight lateral position vs. step width stepping goals without compromising primary task performance.

The same older adults studied here tended to switch off the narrowing paths sooner. This was mediated by their somewhat lower self-perceived balance abilities (Kazanski and Dingwell, 2021). Here, we show that despite their earlier transitions, these older adults retained similar values for 3 of 5 lateral stepping variability and regulation characteristics at switching instances (Figs. 5-6, right). These complimentary analyses thus further illuminate how age-related changes in stepping regulation on these paths still allowed these older adults to achieve appropriate task performance.

In this experiment, we allowed participants to switch from the continuously-narrowing path to the fixed-width path when they chose (Fig. 1). We then analyzed all steps obtained from all participants in each group who walked at each path width (Figs. 5A-C and 6A-B). Because different participants switched paths at different path widths, and because OH participants tended to switch sooner (Fig. 4), the number of steps included in each group-wise bin decreased as path width narrowed, more so for the OH group. This was an inherent limitation of our analyses. Indeed, this was an unavoidable, but also intentional aspect of our study design. A separate aim of this study, as reported in (Kazanski and Dingwell, 2021), was to determine how older adults’ self-perceptions influenced their lane choices. Thus, we intentionally designed these lanes to narrow enough (5 cm) that participants would be compelled to switch before they reached the end.

The present results should be interpreted within the context of our sample population. The OH adults we tested performed both the TUG and FSST tasks significantly slower than YH adults (Table 1). Yet their scores remained well within age-matched ranges for otherwise healthy, non-frail older adults, for both the TUG (Bohannon, 2006; Ibrahim et al., 2017) and FSST (de Aquino et al., 2022; Dite and Temple, 2002). Likewise, these OH adults also reported significantly decreased self-perceptions of their balance abilities (lower ABC and higher I-FES scores) relative to YH (Table 1). Yet their scores again remained far from reported falling concern thresholds, for both the ABC (Lajoie and Gallagher, 2004) and I-FES (Delbaere et al., 2011) assessments.

Together, these findings indicate these OH adults did exhibit actual physical balance deficits, of which they themselves were also aware. Nevertheless, the present study results should be interpreted with caution in that it is not yet clear to what extent these findings might generalize to more frail older adults (de Aquino et al., 2022) and/or other impaired populations.

Older adults often fall following improper or inadequate stepping adjustments, especially on walking paths that change in more challenging environments (Li et al., 2006; Rubenstein, 2006). Here, our study went beyond generic measures of age-related increases in variability magnitude (Brach et al., 2005) to also quantify both the structure of lateral stepping variability and how it connects to underlying multi-objective stepping regulation processes that affect walking balance. We showed that age-related changes in the structure of stepping variability on a continuously-narrowing path reflect step-to-step adaptations in how older adults regulate their movements to promote balance. Our novel implementation of GEM analyses to this distinctly non-straight-walking task demonstrates that walking humans flexibly alter their multi-objective lateral stepping regulation in both age- and context-dependent ways to continuously adjust steps in real time in response to ongoing changes in challenging walking environments.

## ACKNOWLEDGEMENTS

The authors thank Anna C. Render and David M. Desmet for their contributions and technical support throughout data collections. This work was supported by NIH grant 1-R01-AG049735 to JBD & JPC.

## REFERENCES

Arvin, M., Mazaheri, M., Hoozemans, M.J.M., Pijnappels, M., Burger, B.J., Verschueren, S.M.P., van Dieën, J.H., 2016. Effects of narrow base gait on mediolateral balance control in young and older adults. J. Biomech. 49, 1264–1267.

Beauchet, O., Allali, G., Annweiler, C., Bridenbaugh, S., Assal, F., Kressig, R.W., Herrmann, F.R., 2009. Gait Variability Among Healthy Adults: Low and High Stride-to-Stride Variability Are Both a Reflection of Gait Stability. Gerontology 55, 702–706.

Bohannon, R.W., 2006. Reference values for the timed up and go test: a descriptive meta-analysis. J. Geriatr. Phys. Ther. 29, 64–68.

Brach, J.S., Berlin, J.E., VanSwearingen, J.M., Newman, A.B., Studenski, S.A., 2005. Too much or too little step width variability is associated with a fall history in older persons who walk at or near normal gait speed. J. Neuroeng. Rehabil. 2, 21.

Bruijn, S.M., van Dieën, J.H., 2018. Control of human gait stability through foot placement. J. R. Soc. Interface 15, 1–11.

Burns, E.R., Kakara, R., 2018. Deaths from Falls Among Persons Aged ≥65 Years — United States, 2007–2016. Morb. Mortal. Wkly. Rep. 67, 509–514.

Butler, A.A., Lord, S.R., Taylor, J.L., Fitzpatrick, R.C., 2015. Ability Versus Hazard: Risk-Taking and Falls in Older People. The Journals of Gerontology Series A: Biological Sciences and Medical Sciences 70, 628–634.

Christou, E.A., 2011. Aging and Variability of Voluntary Contractions. Exercise and Sport Sciences Reviews 39, 77–84.

Cusumano, J.P., Cesari, P., 2006. Body-Goal Variability Mapping in an Aiming Task. Biol. Cybern. 94, 367–379.

Cusumano, J.P., Dingwell, J.B., 2013. Movement Variability Near Goal Equivalent Manifolds: Fluctuations, Control, and Model-Based Analysis. Hum. Mov. Sci. 32, 899–923.

de Aquino, M.P.M., de Oliveira Cirino, N.T., Lima, C.A., de Miranda Ventura, M., Hill, K., Perracini, M.R., 2022. The Four Square Step Test is a useful mobility tool for discriminating older persons with frailty syndrome. Experimental Gerontology 161, 111699.

Delbaere, K., T. Smith, S., Lord, S.R., 2011. Development and Initial Validation of the Iconographical Falls Efficacy Scale. The Journals of Gerontology Series A: Biological Sciences and Medical Sciences 66A, 674–680.

Desmet, D.M., Cusumano, J.P., Dingwell, J.B., 2022. Adaptive Multi-Objective Control Explains How Humans Make Lateral Maneuvers While Walking. PLoS Comput. Biol. 18, e1010035.

Dingwell, J.B., Cusumano, J.P., 2015. Identifying Stride-To-Stride Control Strategies in Human Treadmill Walking. PLoS ONE 10, e0124879.

Dingwell, J.B., Cusumano, J.P., 2019. Humans Use Multi-Objective Control to Regulate Lateral Foot Placement When Walking. PLoS Comput. Biol. 15, e1006850.

Dingwell, J.B., Cusumano, J.P., Rylander, J.H., Wilken, J.M., 2020. How Persons With Transtibial Amputation Regulate Lateral Stepping While Walking in Laterally Destabilizing Environments. Gait & Posture In Revision.

Dingwell, J.B., John, J., Cusumano, J.P., 2010. Do Humans Optimally Exploit Redundancy to Control Step Variability in Walking? PLoS Comput. Biol. 6, e1000856.

Dingwell, J.B., Salinas, M.M., Cusumano, J.P., 2017. Increased Gait Variability May Not Imply Impaired Stride-To-Stride Control of Walking in Healthy Older Adults. Gait Posture 55, 131–137.

Dite, W., Temple, V.A., 2002. A clinical test of stepping and change of direction to identify multiple falling older adults. Arch. Phys. Med. Rehabil. 83, 1566–1571.

Eckardt, N., Rosenblatt, N.J., 2018. Healthy aging does not impair lower extremity motor flexibility while walking across an uneven surface. Hum. Mov. Sci. 62, 67–80.

Herold, F., Aye, N., Hamacher, D., Schega, L., 2019. Towards the Neuromotor Control Processes of Steady-State and Speed-Matched Treadmill and Overground Walking. Brain Topogr 32, 472–476.

Hollman, J.H., McDade, E.M., Petersen, R.C., 2011. Normative spatiotemporal gait parameters in older adults. Gait Posture 34, 111–118.

Ibrahim, A., Singh, D.K.A., Shahar, S., 2017. ‘Timed Up and Go’ test: Age, gender and cognitive impairment stratified normative values of older adults. PLoS ONE 12, e0185641.

Kazanski, M.E., Cusumano, J.P., Dingwell, J.B., 2020. How Healthy Older Adults Regulate Lateral Stepping While Walking in Laterally Destabilizing Environments. J. Biomech. 104, 109714.

Kazanski, M.E., Dingwell, J.B., 2021. Effects of age, physical and self-perceived balance abilities on lateral stepping adjustments during competing lateral balance tasks. Gait Posture 88, 311–317.

Kuo, A.D., Donelan, J.M., 2010. Dynamic principles of gait and their clinical implications. Phys Ther 90, 157–174.

Lajoie, Y., Gallagher, S.P., 2004. Predicting falls within the elderly community: comparison of postural sway, reaction time, the Berg balance scale and the Activities-specific Balance Confidence (ABC) scale for comparing fallers and non-fallers. Arch. Gerontol. Geriatr. 38, 11–26.

Li, W., Keegan, T.H., Sternfeld, B., Sidney, S., Quesenberry, C.P., Jr., Kelsey, J.L., 2006. Outdoor falls among middle-aged and older adults: a neglected public health problem. Am J Public Health 96, 1192–1200.

Lowry, K.A., Sebastian, K., Perera, S., Van Swearingen, J., Smiley-Oyen, A.L., 2017. Age-Related Differences in Locomotor Strategies During Adaptive Walking. J Mot Behav 49, 435–440.

Maidan, I., Eyal, S., Kurz, I., Geffen, N., Gazit, E., Ravid, L., Giladi, N., Mirelman, A., Hausdorff, J.M., 2018. Age-associated changes in obstacle negotiation strategies: Does size and timing matter? Gait Posture 59, 242–247.

Matthis, J.S., Yates, J.L., Hayhoe, M.M., 2018. Gaze and the Control of Foot Placement When Walking in Natural Terrain. Curr. Biol. 28, 1224–1233.e1225.

Moe-Nilssen, R., Helbostad, J.L., 2005. Interstride trunk acceleration variability but not step width variability can differentiate between fit and frail older adults. Gait Posture 21, 164–170.

Moussaïd, M., Helbing, D., Theraulaz, G., 2011. How simple rules determine pedestrian behavior and crowd disasters. Proc. Natl. Acad. Sci. USA 108, 6884–6888.

Orendurff, M.S., Schoen, J.A., Bernatz, G.C., Segal, A.D., Klute, G.K., 2008. How Humans Walk: Bout Duration, Steps per Bout, and Rest Duration. J. Rehabil. Res. Develop. 45, 1077–1090.

Owings, T.M., Grabiner, M.D., 2004. Variability of step kinematics in young and older adults. Gait Posture 20, 26–29.

Render, A.C., Kazanski, M.E., Cusumano, J.P., Dingwell, J.B., 2021. Walking humans trade off different task goals to regulate lateral stepping. J. Biomech. 119, 110314.

Roerdink, M., Geerse, D.J., Peper, C.E., 2021. ‘Haste makes waste’: The tradeoff between walking speed and target-stepping accuracy. Gait Posture 85, 110–116.

Roos, P.E., Dingwell, J.B., 2010. Influence of Simulated Neuromuscular Noise on Movement Variability and Fall Risk in a 3D Dynamic Walking Model. J. Biomech. 43, 2929–2935.

Rosenblatt, N.J., Hurt, C.P., Latash, M.L., Grabiner, M.D., 2014. An apparent contradiction: increasing variability to achieve greater precision? Exp. Brain Res. 232, 403–413.

Rubenstein, L.Z., 2006. Falls in older people: epidemiology, risk factors and strategies for prevention. Age and Ageing 35, ii37–41.

Schrager, M.A., Kelly, V.E., Price, R., Ferrucci, L., Shumway-Cook, A., 2008. The effects of age on medio-lateral stability during normal and narrow base walking. Gait Posture 28, 466–471.

Skiadopoulos, A., Moore, E.E., Sayles, H.R., Schmid, K.K., Stergiou, N., 2020. Step width variability as a discriminator of age-related gait changes. Journal of NeuroEngineering and Rehabilitation 17, 41.

Sun, R., Cui, C., Shea, J.B., 2017. Aging effect on step adjustments and stability control in visually perturbed gait initiation. Gait Posture 58, 268–273.

Todorov, E., 2004. Optimality principles in sensorimotor control. Nat. Neurosci. 7, 907–915.

Twardzik, E., Duchowny, K., Gallagher, A., Alexander, N., Strasburg, D., Colabianchi, N., Clarke, P., 2019. What features of the built environment matter most for mobility? Using wearable sensors to capture real-time outdoor environment demand on gait performance. Gait Posture 68, 437–442.

Vistamehr, A., Kautz, S.A., Bowden, M.G., Neptune, R.R., 2016. Correlations between measures of dynamic balance in individuals with post-stroke hemiparesis. J Biomech 49, 396–400.

Wang, Y., Srinivasan, M., 2014. Stepping in the direction of the fall: the next foot placement can be predicted from current upper body state in steady-state walking. Biol. Lett. 10, 20140405.

Zeni, J.A., Richards, J.G., Higginson, J.S., 2008. Two simple methods for determining gait events during treadmill and overground walking using kinematic data. Gait Posture 27, 710–714.

Zhang, Y., Smeets, J.B.J., Brenner, E., Verschueren, S., Duysens, J., 2021. Effects of ageing on responses to stepping-target displacements during walking. Eur J Appl Physiol 121, 127–140.

