## Supplement 1 for "How Older Adults Regulate Lateral Stepping on Narrowing Walking Paths"

### SUPPLEMENT # 1 — ADDITIONAL BACKGROUND

#### *Walking On Defined Paths*

When humans walk, they regularly negotiate widely varying contexts (Orendurff et al., 2008), both indoors and out (Matthis et al., 2018). In nearly all of these contexts, humans follow some particular *path* that guides not only where they walk (Moussaïd et al., 2011), but also the lateral stepping movements they make to follow their path (Dingwell and Cusumano, 2019). These paths sometimes have well-defined lateral boundaries (e.g., building hallways, store aisles, etc.), but sometimes not (e.g., Matthis et al., 2018; Moussaïd et al., 2011).

When people walk along a path (e.g., hallway, sidewalk), the lateral boundaries of that path explicitly prescribe a primary walking task: stay on the path. We express this mathematically as (Dingwell and Cusumano, 2019):

$$\forall\, n \in \{1,\ldots,N\}: \qquad -\frac{W_P(x)}{2} < \{z_{Ln}, z_{Rn}\} < +\frac{W_P(x)}{2}, \tag{S1}$$

where for any step, $n$, in a sequence of $N$ consecutive steps, $W_P(x_n)$ defines the path width that can change according the walker's progression along the path (i.e., in the $x$-direction), and $\{z_{Ln}, z_{Rn}\}$ define the lateral left and right foot placements relative to the path center (see Fig. 2A in main paper).

We showed in (Dingwell and Cusumano, 2019) that, for straight paths of constant width [i.e., for $W_P(x) = W \equiv$ Constant in Eq. (S1)], humans do not directly regulate $\{z_{Ln}, z_{Rn}\}$ themselves. Instead, they coordinate $\{z_{Ln}, z_{Rn}\}$ to achieve more-general walking task goals related to whole-body movement. In particular, they try to "maintain step width" by minimizing errors in step width ($w$), and "stay on their path" by minimizing errors in lateral body position ($z_B$) relative to the path center. Multi-objective step-to-step regulation of this form is defined by the goal function:

$$\mathbf{F} = \begin{bmatrix} z_{Bn} - z_B^* \\ w_n - w^* \end{bmatrix}, \tag{S2}$$

where $z_{Bn}$ and $w_n$ are the values of the goal states at step $n$, $z_{Bn}^*$ and $w_n^*$ are their "desired" values, and the task *goal* is to drive $\mathbf{F} \rightarrow 0$ (Cusumano and Dingwell, 2013). Defining lateral body position ($z_B$) as the midpoint between the feet and step width ($w$) as the difference between right and left foot placements (Dingwell and Cusumano, 2019):

$$\begin{aligned} z_{Bn} &= \tfrac{1}{2}\cdot\left(z_{Ln} + z_{Rn}\right) \\ w_n &= z_{Rn} - z_{Ln} \end{aligned}, \tag{S3}$$

steps $\{z_{Ln}, z_{Rn}\}$ are then directly related to the goal states $\{z_{Bn}, w_n\}$ by (Dingwell and Cusumano, 2019):

$$\begin{bmatrix} z_{Bn} \\ w_n \end{bmatrix} = \begin{bmatrix} \tfrac{1}{2} & \tfrac{1}{2} \\ -1 & 1 \end{bmatrix} \begin{bmatrix} z_{Ln} \\ z_{Rn} \end{bmatrix}. \tag{S4}$$

#### *Visualizing Multi-Objective Stepping Dynamics*

We can visualize how steps $\{z_{Ln}, z_{Rn}\}$ directly relate to goal states $\{z_{Bn}, w_n\}$, as defined by Eq. (S4), by plotting them in the $[z_L, z_R]$ plane (Fig. S1-1) (Desmet et al., 2022; Dingwell and Cusumano, 2019). Individual goals to maintain constant $z_{Bn} = z_B^*$ and $w_n = w^*$ each define linear Goal-Equivalent Manifolds, or GEMs (Cusumano and Dingwell,

2013), that are orthogonal to each other in this plane (Fig. S1-1A). Each GEM contains infinite possible $\{z_{Ln}, z_{Rn}\}$ combinations that achieve either the target lateral body position, $z_{Bn} = z_B^*$, or target step width, $w_n = w^*$, respectively.

When walking straight ahead, people strongly prioritize maintaining $w$ far more than $z_B$ (Dingwell and Cusumano, 2019). This yields stepping data that are strongly aligned with the $w^*$ GEM (Fig. S1-1A). To enact lateral side-stepping maneuvers, people trade off stability for maneuverability (Desmet et al., 2022). This yields steps that are distributed approximately isotropically in $[z_L, z_R]$ (Fig. S1-1B). If people prioritize maintaining $z_B$ more than $w$, we expect these distributions to become strongly aligned to the $z_B^*$ GEM (Fig. S1-1C) (Dingwell and Cusumano, 2019).

The nature and degree to which people regulate their step placements while walking can then be quantified by fitting 95% prediction ellipses to these $[z_L, z_R]$ plane data clouds (Desmet et al., 2022). The sizes and shapes of these ellipses are then completely defined by their area, aspect ratio ($\lambda_1/\lambda_2$), and angular orientation ($\Delta\theta$), which we measure counter-clockwise relative to the $+w^*$ axis (Fig. S1-1C).

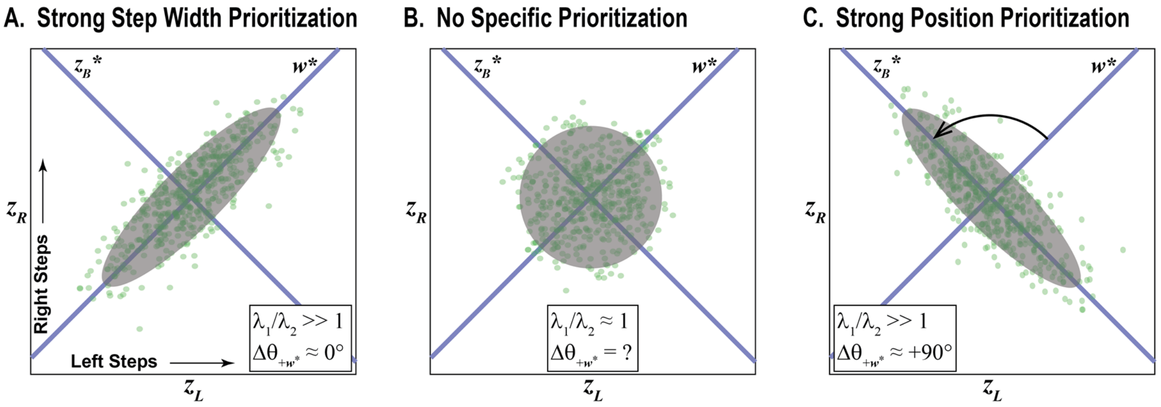


**Figure S1-1: Schematic Depictions of Different Types of Stepping Regulation.** Every step involves both a left ($z_L$) and right ($z_R$) foot placement. Thus, each step yields a single point in the $[z_L, z_R]$ plane. **A:** When over many steps, people strongly prioritize maintaining step width ($w$), these steps will form a cloud of points very strongly aligned to the $\boldsymbol{w}^*$ GEM. **B:** If people prioritize all stepping options roughly equally (such as when they need to be maneuverable), steps will be distributed roughly isotropically (i.e., in the shape of a circle). **C:** If people prioritize maintaining lateral position ($z_B$), such as when walking on a very narrow path, we expect steps to form a cloud of points very strongly aligned to the $\boldsymbol{z_B}^*$ GEM. Corresponding values of the fitted ellipse aspect ratio ($\lambda_1/\lambda_2$) and orientation ($\Delta\theta$) are shown for each case. Note that for data that form a *perfect* circle ($\lambda_1/\lambda_2 = 1$), the orientation angle is undefined.

### REFERENCES

Cusumano, J.P., Dingwell, J.B., 2013. Movement Variability Near Goal Equivalent Manifolds: Fluctuations, Control, and Model-Based Analysis. Hum. Mov. Sci. 32, 899-923.

Desmet, D.M., Cusumano, J.P., Dingwell, J.B., 2022. Adaptive Multi-Objective Control Explains How Humans Make Lateral Maneuvers While Walking. PLoS Comput. Biol. 18, e1010035.

Dingwell, J.B., Cusumano, J.P., 2019. Humans Use Multi-Objective Control to Regulate Lateral Foot Placement When Walking. PLoS Comput. Biol. 15, e1006850.

Matthis, J.S., Yates, J.L., Hayhoe, M.M., 2018. Gaze and the Control of Foot Placement When Walking in Natural Terrain. Curr. Biol. 28, 1224-1233.e1225.

Moussaïd, M., Helbing, D., Theraulaz, G., 2011. How simple rules determine pedestrian behavior and crowd disasters. Proc. Natl. Acad. Sci. USA 108, 6884-6888.

Orendurff, M.S., Schoen, J.A., Bernatz, G.C., Segal, A.D., Klute, G.K., 2008. How Humans Walk: Bout Duration, Steps per Bout, and Rest Duration. J. Rehabil. Res. Develop. 45, 1077-1090.
