## Supplement 2 for "How Older Adults Regulate Lateral Stepping on Narrowing Walking Paths"

### SUPPLEMENT # 2 — ADDITIONAL RESULTS

#### *Example Stepping Ellipses*

Figure S2-1 presents YH and OH lateral stepping variability, visualized in [$z_L$, $z_R$] at several individual bin widths ($W_P$). As paths narrowed, both groups remained mostly within the path bounds, but progressively used more of the available path width to vary their lateral steps. In Fig. S2-1, the fitted ellipses fill increasingly larger portions of the available path (gray boxes) as the path narrow (i.e., as $W_P$ decreases from right to left).

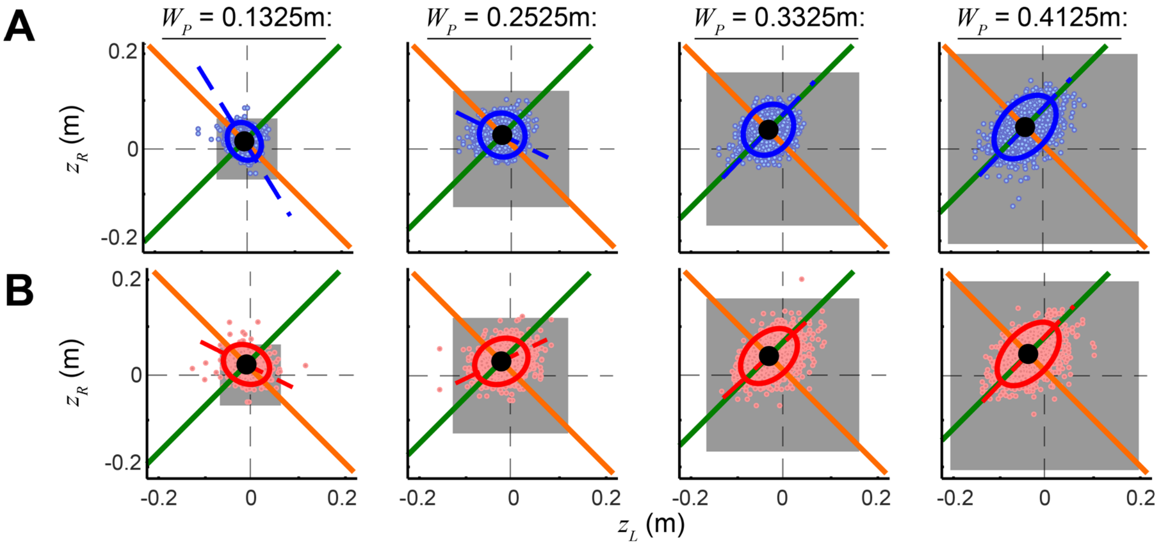


**Figure S2-1: Example Stepping Ellipses. A:** Group-wise lateral stepping data in the [$z_L$, $z_R$] plane for YH adults (blue). **B:** Corresponding [$z_L$, $z_R$] plane data for OH adults (red). Each subplot shows distributions of all lateral steps (individual points) for a single group-wise sub-sample at the designated bin width, $W_P$. Grey squares of area $W_P^2$ indicate the allowable areas where steps would remain inside the designated path bounds. Fitted 95% prediction ellipses and their major axes are shown for each distribution. Diagonal orange lines indicate the constant position ($z_B^*$) GEM. Diagonal green lines indicate the constant step width ($w^*$) GEM.

---

#### *Example Direct Control Plots*

Figures S2-2 & S2-3 present YH and OH error correction of $z_B$ and $w$, respectively, at several individual bin widths ($W_P$). Initially at wider $W_P$, both groups exhibited strong correction of $w$ errors (i.e., $M(w) \rightarrow -1$) and weak correction of $z_B$ errors (i.e., $M(z_B) \rightarrow 0$), consistent with the prioritization observed during continuous, straight-ahead walking (Dingwell and Cusumano, 2019). As paths narrowed, both age groups decreased their $w$ error correction while increasing their $z_B$ error correction.

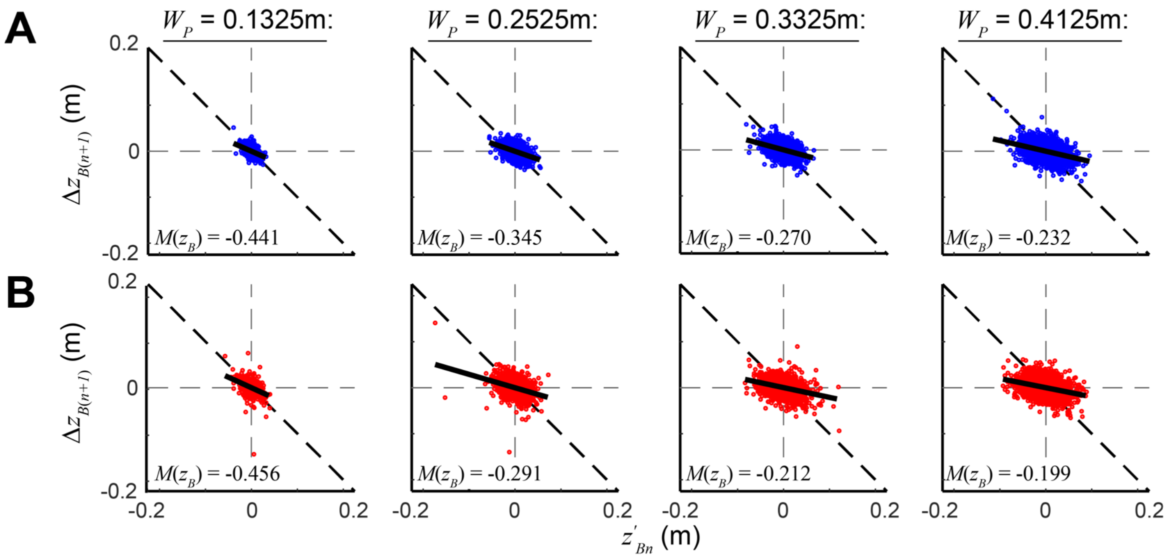


**Figure S2-2: Example Direct Control Plots – Position Control.** **A:** Example step-to-step error correction (Fig. 3C) of lateral body position ($z_B$) for YH adults (blue). **B:** Corresponding step-to-step error correction of $z_B$ for OH adults (red). Each subplot shows $z_B$ correction for a single group-wise sub-sample at the designated bin width, $W_P$. Diagonal solid black lines indicate least-squares fits for each set of data points. Slopes of these lines, $M(z_B)$, are shown on each respective plot. These slopes quantify the degree of direct $z_B$ error correction from step-to-step within that sub-sample.

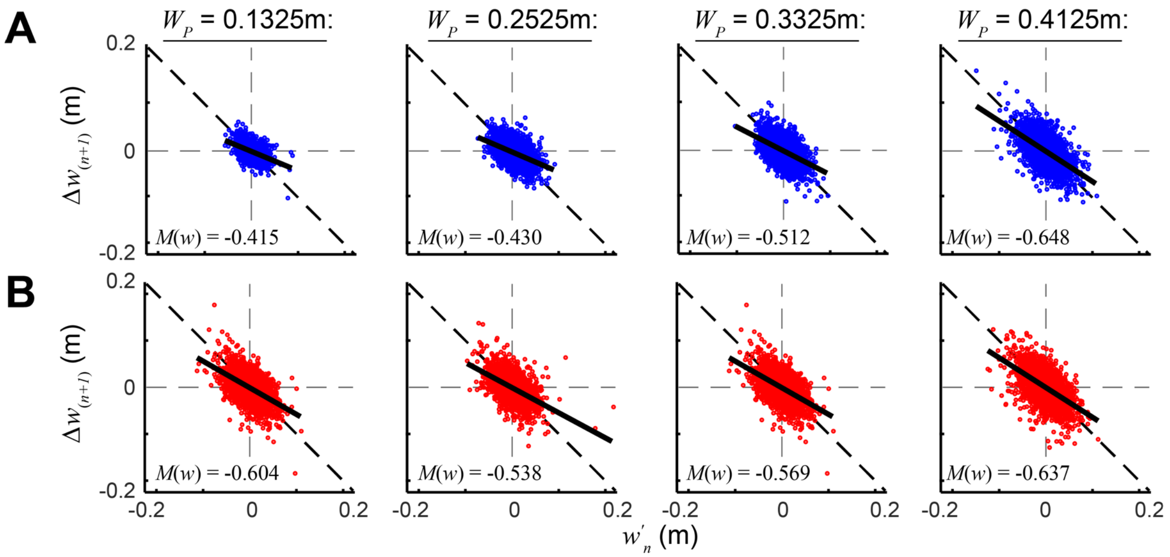


**Figure S2-3: Example Direct Control Plots – Step Width Control.** **A:** Example step-to-step error correction (Fig. 3C) of step width ($w$) for YH adults (blue). **B:** Corresponding step-to-step error correction of $w$ for OH adults (red). Each subplot shows $w$ correction for a single group-wise sub-sample at the designated bin width, $W_P$. Diagonal solid black lines indicate least-squares fits for each set of data points. Slopes of these lines, $M(w)$, are shown on each respective plot. These slopes quantify the degree of direct $w$ error correction from step-to-step within that sub-sample.
